# Blood-brain barrier model architecture shapes peripheral immune cell trafficking in Parkinson’s disease

**DOI:** 10.64898/2026.06.04.730214

**Authors:** Oceane Martinez, Lucile Gentile, Floriane Bretheau, Allison Yearwood, Oceane Arevalo, Flavia Natale Alves Martins Borba, Martine Boutin, Eric Boilard, Kelvin Luk, Francesca Cicchetti, Aurelie de Rus Jacquet

## Abstract

Parkinson’s disease (PD) is a neurodegenerative disorder traditionally characterized by dopaminergic neuron loss in the *substantial nigra pars compacta*, but peripheral immune dysregulation and blood-brain barrier (BBB) dysfunction have been increasingly implicated in disease etiology. However, how circulating immune cells interact with the human BBB and how these interactions are captured across experimental models remains poorly understood. In particular, human BBB models offer multiple platforms to interrogate these biological questions, with organ-on-chip approaches attracting significant interest. In this context, it is essential to determine whether model architecture influences the assessment of immune-endothelial interactions in PD, and if it may lead to fundamentally different interpretations of immune cell trafficking at the human BBB. Here, we compared peripheral blood mononuclear cells (PBMCs) from control donors or individuals with PD in human induced pluripotent stem cell (iPSC)-derived BBB models to determine how static and dynamic BBB systems influence immune cell behavior. To do so, we leveraged our brain chip platform to establish a model based on expression of the PD-associated *SNCA* triplication mutation. Using a two-dimensional transwell system and a three-dimensional (3D) microfluidic BBB chip, we evaluated PBMC attachment and transmigration under conditions of PBMC disease status, endothelial genotype associated with *SNCA* triplication, and exposure to α-synuclein (α-Syn) monomers or preformed fibrils (PFFs). PBMCs from PD donors showed increased baseline reactivity and altered endothelial interactions compared with controls. In transwell models, *SNCA* triplication increased PBMC attachment and selectively enhanced PD PBMC transmigration, while PFF increased attachment without affecting transmigration. In contrast, in the microfluidic BBB chip, attachment was largely unchanged by endothelial genotype or α-Syn exposure, whereas transmigration increased following α-Syn monomer pre-treatment. Together, PBMC-BBB interactions in PD appear to be shaped by immune cell status, endothelial genotype, and α-Syn exposure, but are strongly influenced by BBB model dimensionality and flow. This study underscores the importance of physiologically relevant multicellular and flow-based BBB systems and provides a human-focused framework for studying peripheral immune cell trafficking across the diseased BBB. These findings also emphasize that biological insights into BBB function are inherently shaped by the experimental model used, underscoring the need for complementary human BBB platforms.

## Introduction

Parkinson’s disease (PD) is a progressive neurodegenerative disorder characterized by the selective loss of dopaminergic neurons in the *substantia nigra pars compacta* and the accumulation of misfolded α-synuclein (α-Syn) into Lewy Bodies and Lewy Neurites. Beyond neuronal loss, clinical and experimental evidence supports a contribution of non-neuronal processes to PD pathology, including peripheral immune activation and neurovascular alterations ^1–4^. While most cases are idiopathic, rare genetic forms have provided critical insights into disease mechanisms. Notably, genomic triplication of *SNCA*, the gene encoding α-Syn, causes a severe, early-onset form of PD, demonstrating that increased α-Syn gene dosage is sufficient to drive neurodegeneration. These findings establish α-Syn dysregulation as a central pathogenic event in PD and underscore the causal link between protein overexpression, misfolding, and dopaminergic neuron vulnerability ^5–7^. Studies have suggested that α-Syn preformed fibrils (PFFs) can trigger robust inflammatory responses in peripheral blood mononuclear cells (PBMCs), as demonstrated by increased cytokine production by PD and control PBMCs ^8^. In addition, α-Syn PFFs have been reported to activate innate immune signaling pathways and promote pro-inflammatory gene expression in monocytes ^9^. These immune responses correlate with disease severity, suggesting that pathological forms of α-Syn may play a role in PD progression through peripheral immune modulation ^10^. In line with this concept, T cells from individuals with PD can recognize α-Syn-derived peptides, indicating that adaptive immune responses may also contribute to disease pathology ^11–13^.

In this context, the peripheral-brain immune axis is increasingly recognized as a key component of PD pathophysiology. Immune cell trafficking across the blood-brain barrier (BBB) is tightly regulated under physiological conditions to protect the central nervous system (CNS) from peripheral inflammation ^14,15^. The BBB consists primarily of brain microvascular endothelial cells (BMECs), astrocytes, and pericytes, forming a dynamic multicellular structure that regulates molecular fluxes and immune cell entry into the CNS ^16^. Beyond its barrier function, the BBB actively controls immune cell adhesion and transmigration, and disease-associated alterations in BMEC phenotype or barrier integrity can dysregulate immune cell trafficking ^17^. Evidence indicates that peripheral immune cells can infiltrate the CNS in PD, highlighting the importance of BBB regulation in this process. Postmortem analyses have revealed increased numbers of CD4+ and CD8+ T lymphocytes in regions of dopaminergic neurodegeneration, including the *substantia nigra*, indicating recruitment of adaptive immune cells from the periphery ^18–20^. In parallel, markers associated with monocyte-derived cells have been detected in *in vivo* PD models ^21,22^, suggesting that peripheral myeloid populations may also contribute to neuroinflammatory processes, although distinguishing infiltrating monocytes from resident microglia in human remains challenging. Experimental *in vivo* PD models further support the ability of peripheral T cells and monocytes to enter the brain and exacerbate dopaminergic dysfunction ^18,23^. Together, these findings support a model in which peripheral immune activation and BBB dysregulation converge to promote immune cell infiltration into the PD brain, linking systemic immune alterations to central neurodegenerative processes. Despite increasing *in vivo* evidence from animal models supporting peripheral immune involvement and BBB dysfunction in PD ^24^, a detailed understanding of these processes in human-relevant systems remains limited. Accordingly, direct interrogation of immune cell interactions with the human BBB relies on *in vitro* models, which represent the principal translational platform for studying these interactions. However, it remains unclear whether circulating immune cells interact with the human BBB in a manner that is consistently captured across experimental BBB platforms.

Novel human-focused *in vitro* models may prove useful to study immune-BBB interactions. Transwell systems, widely used since the 1990s, culture BMECs on porous membranes and provide a simple and standardized platform ^16^. However, they lack physiological shear stress and multicellular complexity, limiting their ability to reproduce dynamic BBB functions such as immune cell adhesion and transmigration, thereby reducing translational relevance ^16,25,26^. To overcome these limitations, microfluidic BBB chips have been developed to better recapitulate the 3D and dynamic nature of the BBB, and more accurately capture complex interactions and investigate immune cell infiltration ^27,28^. These systems incorporate controlled fluid shear stress, enabling physiologically relevant endothelial organization and barrier function ^27–29^. They also allow co-culture of BMECs with CNS cell types under flow, enabling more biomimetic interactions between cells forming the neurovascular unit (NVU) ^29^, and improving modeling of neuroinflammatory responses and immune cell extravasation ^27^. Static transwell systems and dynamic microfluidic platforms differ substantially in dimensionality, cellular complexity, and exposure to physiological shear stress, all of which may influence immune cell interactions. Importantly, whether static and dynamic human BBB models generate convergent or divergent interpretations of immune cell trafficking behaviors remains largely unexplored. Addressing this question is particularly relevant in PD, where both peripheral immune activation and BBB dysfunction are implicated in disease progression.

In this study, we used human induced pluripotent stem cell (iPSC)-derived BBB models to investigate how peripheral immune cell interactions with the BBB are modulated in PD. By combining transwell and microfluidic BBB platforms with PBMCs from control donors and individuals with PD, we evaluated immune cell attachment, transmigration, and inflammatory responses under distinct experimental conditions. We further examined the influence of endothelial genotype associated with *SNCA* triplication and pre-exposure to α-Syn monomers or PFFs. This approach enables direct comparison of static and dynamic BBB systems to determine whether model architecture leads to fundamentally different interpretations of immune cell trafficking at the human BBB.

## Material and Methods

### Isolation of PBMCs

Blood samples were collected from healthy and PD volunteers following informed consent, under ethical approval number A13-02-1096 by the Comité d’éthique de la recherche du CHU de Québec. Blood was collected into heparin-coated tubes and processed on the day of collection. PBMCs were isolated using SepMate tubes (Stemcell Technologies, #85450) in combination with LymphoPrep density gradient medium (Stemcell Technologies, #85450). Briefly, whole blood was diluted to a final volume of 20 mL with phosphate-buffered saline (PBS) supplemented with 2% fetal bovine serum (FBS). Fifteen mL of LymphoPrep were added to SepMate tubes, and the diluted blood was carefully layered on top of the density gradient. Samples were centrifuged at 1200 x g for 10 minutes (min) at room temperature, according to the manufacturer’s instructions. Following centrifugation, the upper compartement of the SepMate tube containing the PBMC-enriched fraction was poured into a 50 mL conical tube and washed with PBS containing 2% FBS. Cells were centrifuged at 300 x g for 8 min at room temperature. The resulting cell pellet was resuspended in freezing medium consisting of FBS supplemented with 10% dimethyl sulfoxide (DMSO) and cryopreserved until further use.

### Culture and differentiation of iPSCs

#### iPSCs lines and maintenance

Two female human iPSC lines were used in this study. The control iPSC line was kindly provided by Prof. Dr. Thomas Gasser (Universitätsklinikum Tübingen) and Prof. Dr. Hans R. Schöler (Max-Planck Institute) and has been previously described ^30^. The *SNCA* triplication iPSC line was a generous gift from Dr. Randall Moon (Howard Hughes Medical Institute/University of Washington); the original fibroblasts were obtained from the Coriell Institute for Medical Research via the NINDS repository (ND 27760). Human iPSCs were maintained in mTeSR Plus medium (StemCell Technologies, Vancouver, BC, Canada, #100-0276), cultured on Geltrex-coated dishes (Thermo Fisher Scientific, Waltham, MA, # A1413301), and passaged as small aggregates using ReLeSR (StemCell Technologies, #5872), according to the manufacturer’s instructions.

#### Differentiation of iPSCs into astrocytes and dopaminergic neurons

Differentiation of iPSCs into astrocytes was performed as previously described and extensively characterized ^29,31,32^. Briefly, iPSCs were first neuralized by dual SMAD inhibition to generate midbrain-patterned neural progenitor cells (NPCs). NPCs were subsequently differentiated into astrocytes through serial passaging and culture in astrocyte medium (ScienCell Research Laboratories, #1801) for a total duration of four weeks. Newly differentiated astrocytes were either cryopreserved for subsequent experiments or plated directly for downstream assays.

To generate dopaminergic neurons, NPCs were plated at 7 × 10^6^ cells per well on a Geltrex-coated 6-well plates. Twenty-four hours (h) after plating, cells were rinsed with Dulbecco’s PBS (DPBS) and cultured in neuron differentiation medium consisting of Neurobasal medium supplemented with 1x B-27 (without vitamin A; FisherScientic, #12587010), 0.5 mM dibutyryl cAMP, 10 µM DAPT, 0.2 mM ascorbic acid, 20 ng/mL brain-derived neurotrophic factor (BDNF; ThermoFisher Scientific, #450-02), 20 ng/mL of glial cell line-derived neurotrophic factor (GDNF; ThermoFisher,#450-10) ^32,33^. The differentiation medium was refreshed every 2 days for a total differentiation period of 7-10 days. As reported previously, the resulting cultures consist of approximately 70% dopaminergic neurons, with the remaining cells comprising non-dopaminergic neuronal populations.

#### Differentiation of iPSCs into BMEC-like cells

Differentiation of iPSCs into BMEC-like cells was performed following established protocols, with minor modifications (Lippmann et al., 2012; Hollmann et al., 2017). iPSCs were maintained in TeSR^TM^-E8 medium (StemCell Technologies, #05990) prior to differentiation. For BMEC induction, iPSCs were dissociated into single cells using Accutase (Millipore Sigma, #A6964) and seeded at a density of 12,500 cells/cm² onto Geltrex-coated plates in E8 medium supplemented with ROCK inhibitor Y-27632 (StemCell Technologies, #72302) (day -1). After 24 h, cultures were transitioned to TeSR-E6 medium (StemCell Technologies, #05946), which was refreshed daily for four consecutive days (days 0–3). On days 4 and 5, the medium was replaced with endothelial cell (EC) medium consisting of human endothelial serum-free medium (Thermo Fisher Scientific, #11111044) supplemented with 1% platelet-poor human plasma (PPP; Sigma, #P2918), 20 ng/mL basic fibroblast growth factor (FGF-basic; ThermoFisher Scientific, #100-18C), and 10 µM retinoic acid (Sigma, #R2625). On day 6, BMEC-like cells were detached using Accutase and replated onto Geltrex-coated dishes or directly into microfluidic chips for subsequent experiments, using EC medium supplemented with 20 µM RO-20-1724, 400 µM dibutyryl cAMP, and 10 µM retinoic acid (referred to as EC^+^ medium), and 10 µM Y-27632 (Selleck Chemicals, #S1049).

### Preparation of α-Syn monomers and PFFs

Wildtype α-Syn protein production, purification, and fibrilization were conducted as previously described ^34^. Briefly, full-length human α-Syn cDNA was expressed in E. coli BL21 (DE3) RIL cells. Following homogenization, bacterial lysates were boiled and then separated using size exclusion (Superdex 200; Cytiva) and anion exchange (HiTrapQ HP; Cytiva) chromatography. Monomeric protein was further processed with endotoxin removal spin columns (Pierce) until LPS levels reached <1 EU/mg protein as measured using a Limulus amebocyte lysate (LAL) endotoxin quantification kit (Pierce). PFF assembly was achieved by diluting the purified α-Syn to 5 mg/ml (360 µM) in sterile DPBS (without Ca^2+^/Mg^2+^pH 7.0; Mediatech) and then agitated constantly at 1,000 rpm at 37°C for 7 days. Reaction completion was confirmed by ultracentrifugation, thioflavin T fluorescence, and activity in primary neuronal cultures prepared from CD1 mouse embryos (E16-18) as previously described ^35^.

### PBMCs treatment with α-Syn monomers and PFFs

Cryopreserved PBMCs were rapidly thawed at 37°C, when indicated, labelled with CellBrite Green membrane dye (1:200 dilution; Biotium #30021) for 15 min at 37°C. Cells were washed twice with RPMI 1640 basal medium (Thermo Fisher Scientific, #11875093) to remove excess dye and plated in individual wells of 48-well plates at a density of 1 x 10^6^ cells/cm^2^. PBMCs used for transwell migration assays were not labelled with CellBrite to avoid interference with luminescence-based readouts. PBMC culture medium consisted of RPMI 1640 supplemented with 5% FBS, 1% GlutaMAX^TM^, and 1% Penicillin-Streptomycin. Twenty-four hours after plating, PBMC medium was supplemented with α-Syn monomers, PFFs, or vehicle control (PBS) to a final concentration of 5 µg/mL. Prior to treatment, PFFs preparations were fragmented by sonication in a water bath sonicator (Branson 1510 model) for 5 min in cold water. PBMC cultures were maintained for a total duration of 7 days, with medium replenished every 2-3 days using fresh PBMC medium containing the corresponding α-Syn species or vehicle control.

### PBMCs attachment to BMEC-like cell monolayers

BMEC-like cells differentiated from iPSCs were detached using Accutase and plated onto Geltrex-coated black, clear bottom 96-well plates at a density of 35,000 cells per well. Cells were allowed to form a confluent monolayer overnight. Control and PD PBMCs pre-exposed for 7 days to α-Syn monomers, PFFs, or vehicle control were harvested, washed with PBS, and incubated in PBMC medium supplemented with 0.5 µg/mL calcein AM for 15 min at 37 °C. In parallel, BMEC-like cell monolayers were incubated with Hoechst nuclear stain for 15 min at 37°C to visualize endothelial nuclei. Following staining, PBMCs were washed twice with DPBS to remove unbound dye. Labelled PBMCs were then added to BMEC-like cell monolayers at a density of 20,000 PBMCs per well and incubated for 15 min at 37°C in a humified cell culture incubator. After incubation, non-adherent PBMCs were removed by gentle washing with DPBS. Image acquisition was performed using a Cytation 5 Cell Imaging Multi-Mode Reader (BioTek, Agilent Technologies). The entire surface of each well was imaged. Images were processed and analyzed using Fiji software (version 1.53T). Background subtraction was applied independently to each fluorescence channel (green: calcein AM; blue: Hoechst), followed by thresholding to identify PBMCs (calcein AM) and BMEC-like cell nuclei (Hoechst). The number of adherent PBMCs and endothelial nuclei was quantified using the *Analyze Particles* tool.

### PBMCs migration in the 2D transwell BBB model

BMEC-like cells differentiated from iPSCs were detached using Accutase and seeded onto Geltrex-coated Transwell inserts (6.5 mm diameter, 5 µm pore size; Corning Transwells, Sigma #CLS3421) at a density of 35,000 cells per insert. Cells were plated in 100 µl EC^+^ medium supplemented with 10 µM ROCK inhibitor. To prevent dehydration of the inserts, 600 µl of EC^+^ medium was added to the lower compartment. BMEC-like cells were allowed to adhere and form monolayers overnight. Control and PD PBMCs pre-exposed for 7 days to α-Syn monomers, PFFs, or vehicle control were harvested, washed with DPBS, and added to the upper compartment of the inserts at a density of 3.75 x 10^4^ PBMCs per insert (100 µL volume, PBMC medium). To establish a chemotactic gradient, the lower compartment was supplemented with recombinant human interleukin-6 (IL-6; 50 ng/mL) and interleukin-8 (IL-8; 50 ng/mL) diluted in EC^+^ medium (600 µL total volume), as previously described to model glial-derived immune signaling. After overnight incubation at 37°C, the plates were removed from the incubator and allowed to equilibrate at room temperature for 20 min. Transwell inserts were then carefully removed, and the entire volume of medium from the lower compartment was collected and mixed thoroughly with CellTiter-Glo Luminescent Cell Viability Assay Kit (Promega, #G7572) at a 1:1 ratio, according to the manufacturer’s instructions. Following a 10 min incubation in the dark, luminescence was measured using a microplate reader (BioTek, Synergy, Agilent Technologies) to quantify migrated PBMCs.

### Preparation of 3D microfluidic BBB chips

#### BBB chip platform

BBB chips were generated using the OrganoPlate 3-lane 40 platform (Mimetas, Gaithersburg, MD), as previously described ^29,36^. Each microfluidic chip comprises three parallel channels: a top channel for formation of the endothelial-like vessel, a central channel containing a collagen I extracellular matrix, and a bottom channel representing the CNS compartment. The vascular channel is seeded with iPSC-derived BMEC-like cells, and the CNS channel is seeded with primary human brain vascular pericytes (ScienCell Research Laboratories, #1200), iPSC-derived astrocytes, and iPSC-derived dopaminergic neurons.

#### Extracellular matrix preparation and channel coating

One day prior to cell seeding, the central channel was filled with 1.8 μL of collagen I solution (7.5 mg/mL; Selleck Chemicals, #CLS354249-1EA) and incubated at 37°C for 10-15 min. To prevent dehydration, 50 μL of Hank’s Balanced Salt Solution (HBSS) was added to the adjacent reservoirs. The collagen matrix was allowed to polymerize for an additional 60 min at 37°C. Subsequently, the vascular channel was coated with 1.8 μL of a fibronectin (50 μg/mL; Sciencell, #8248) and collagen IV (330 μg/mL; Millipore Sigma, #AB769) solution and maintained in the cell culture incubator until cell loading.

#### Cell preparation and loading

iPSC-derived BMEC-like cells were rinsed with DPBS and enzymatically detached using Accutase for 20 min at 37°C. Cells were collected by centrifugation (180 × g for 5 min) and resuspended in EC^+^ medium supplemented with 10 µM ROCK inhibitor. Human primary pericytes were washed with DPBS, briefly exposed to 0.25% trypsin–EDTA for 1 min, pelleted (180 × g for 5 min), and resuspended in astrocyte medium. iPSC-derived astrocytes were either rapidly thawed from cryostorage immediately prior to use or detached from maintenance cultures using Accutase for 5 min, followed by centrifugation (180 × g, 5 min) and resuspension in astrocyte medium. For microfluidic loading, BMEC-like cells were adjusted to a density of 7 × 10⁴ cells/µL, and 2 µL of the suspension were introduced into the upper-left inlet of each chip using an electronic single-channel pipette. A second mixed suspension containing astrocytes (6 × 10³ cells/µL), pericytes (6 × 10³ cells/µL), and dopaminergic neurons (5 × 10³ cells/µL) was prepared, and 2 µL were delivered into the lower-left inlet. Following cell seeding, 50 µL of EC^+^ medium supplemented with 10 µM ROCK inhibitor was added to the upper-left inlet, while 50 µL of astrocyte medium was added to the lower-left inlet. The OrganoPlate was positioned upright using the manufacturer-provided plate holder and incubated at 37°C for 3-4 h to allow cell attachment. Thereafter, all inlets and outlets were replenished with astrocyte medium in the CNS compartment and EC^+^ medium supplemented with ROCK inhibitor and 3 µM CHIR99021 in the vascular compartment. The OrganoPlate was placed on a perfusion rocker (Mimetas) with a 7° inclination angle alternating every 8 min.

#### Barrier integrity assay

Four days after BBB chip assembly, barrier integrity was assessed as previously described ^29,36^. Briefly, the vascular compartment was perfused with EC medium containing either 3 kDa dextran-Alexa Fluor 680 (12.5 µg/mL; Millipore Sigma,# D34681) or rhodamine B (250 µg/mL; Millipore Sigma) by adding 40 µL to the top-left inlet and 30 µL to the top-right outlet. In parallel, CNS inlets and outlets were replaced with 20 µL of fresh EC medium for the duration of the assay. Live imaging was initiated immediately using a Cytation 5 Cell Imaging Multi-Mode Reader (BioTek, Agilent Technologies. Six images were acquired per chip, with a minimum interval of 6 min between acquisitions. The field of view was adjusted to simultaneously capture the vascular, extracellular matrix, and brain compartments. Fluorescence intensity corresponding to tracer diffusion from the vascular to the brain compartment was quantified using Fiji software (version 1.53T), and apparent permeability (P_AAP_) values were calculated using the following formula:

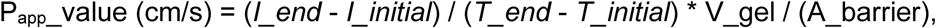

where cm = centimeter, s = seconds; I_intial = initial intensity; I_end = endpoint intensity; T_initial = time intial in seconds; T_end = time end in seconds; V_gel = 1.04 x 10^4^, and A_barrier = 5.7 x 10^3^. A_Barrier and V_gel were considered constants since the area available for molecule exchange was constrained by the dimensions of the plate. In these conditions, the assumption is that the fluid exchange area between the vessel and the brain compartments remains constant regardless of vessel width. P_AAP_ values were used to identify functional BBB chips prior to PBMC perfusion into the vessels, and non-functional chips were excluded from subsequent analyses.

#### PBMCs perfusion in BBB chips

Four days after BBB assembly and following validation of barrier integrity, functional microvessels were selected for PBMC perfusion. Fluorescently labelled PBMCs were harvested, counted, and resuspended in PBMC medium. A total of 45,000 PBMCs were prepared in 100 µL PBMC medium, and 50 µL of the suspension were added to both the inlet and outlet of the vascular channel of each BBB chip. The OrganoPlate was then returned to the cell culture incubator and placed on the perfusion rocker until downstream analysis.

### PBMCs attachment and migration in BBB chips

After a total of 3 days PBMC perfusion in the 3D microfluidic vessels, BBB chips were washed three times with DPBS and fixed with 4% paraformaldehyde (PFA; Fisher Scientific, Waltham, MA) for 20 min at room temperature. The number of fluorescently labelled PBMCs present in the vascular and CNS compartments was subsequently quantified by confocal microscopy using a 20x objective. Quantifications were performed manually by an experimenter blinded to the experimental conditions for the duration of the analysis.

### ELISA

Conditioned media were collected from BBB microfluidic devices or PBMC cultures and centrifuged at 2,000 × g for 10 min to remove cellular debris. Supernatants were aliquoted into single-use samples and stored at -80°C until analysis. Prior to cytokine quantification, aliquots were thawed on ice. Concentrations of IL-6, IL-8, C-X-C motif chemokine ligand 10 (CXCL10), and tumor necrosis factor-α (TNFα) were quantified using commercially available ELISA kits according the manufacturers’ instructions (IL-6, R&D Systems, cat # DY206; IL-8, R&D Systems, cat # DY208; CXCL10, R&D Systems, cat # DY266-05; TNF-α, R&D Systems, cat # DY210-05).

### Immunofluorescence

#### Immunofluorescence on cell monolayers

Cell monolayers were cultured on Geltrex-coated German glass coverslips (Electron Microscopy Sciences, Hatfield, PA), washed once with DPBS, and fixed with 4% PFA for 20 min at room temperature. Following fixation, cells were washed once with DPBS to remove residual PFA, then permeabilized and blocked for 1 h at room temperature in blocking buffer containing 10% (v/v) FBS, 1% (w/v) bovine serum albumin (BSA; Cytiva, #SH30574.02) and 0.3% (v/v) Triton-X100 (Sigma Aldrich, #X100-100ML) diluted in DPBS. Samples were incubated overnight at 4°C with primary antibodies diluted in antibody incubation buffer (1% (w/v) BSA in DPBS). The following primary antibody were used: anti-GFAP (1:200; R&Dystems, #MAB25941), anti-Vimentin (1:500; R&D Systems, #MAB2105), anti-CD44 (1:500; BD Biosciences, #550538), anti-VE-cadherin (1:500; R&D, #AF938), anti-Claudin-5 (1:500; Thermo Fisher Scientific, #35-2500), anti-Zonula occludens-1 (ZO-1; 1:500; Invitrogen, #61-7300), anti-GLUT-1 (1:500; Novus, #NBP2-75785), anti-TUJ-1 (1:500; Millipore Sigma, MAB1637), anti-TH (1:500; PhosphoSolutions, #2025-THRAB), anti-PDGFRβ (1:100; Abcam, #ab32570), and anti-NG2 (1:500; Cell Signalling Technology, #4391). The following day, samples were washed twice with DPBS and incubated for 1 h at room temperature with Alexa Fluor-conjugated secondary antibodies (donkey anti-mouse, donkey anti-rabbit, or donkey anti-goat; Alexa Fluor 488, 555, or 647; Thermo Fisher Scientific) diluted 1:1,000 in antibody incubation buffer. After two DPBS washes, coverslips were incubated with DAPI (1:5,000 dilution) for 10 min, followed by three additional DPBS washes and mounting onto glass slides using ProLong Diamond Antifade. Slides were cured for 24 h at room temperature in the dark prior to imaging.

#### Immunofluorescence in BBB microfluidic chip

BBB chips were fixed in 4% PFA for 20 min at room temperature and washed three times with DPBS. After removal of DPBS from the channels, a permeabilization and blocking solution containing 0.1% (w/v) saponin, 1% (v/v) FBS, and 10% (w/v) BSA in PBS was added to the inlets and outlets. Chips were incubated for 1 h at room temperature in the dark. Following blocking, microfluidic channels were washed twice with DPBS and incubated overnight at 4°C with primary antibodies diluted 1:50 in antibody incubation buffer. The following day, primary antibody solutions were removed and the chips were washed twice with DPBS before incubation with Alexa Fluor-conjugated secondary antibodies (1:500 in antibody incubation solution), for 1 h at room temperature in the dark. After secondary antibody incubation, chips were washed three times with DPBS and incubated with Alexa Fluor 546 phalloidin (1:1000; Invitrogen, #A22283) in antibody incubation buffer for 20 min at room temperature in the dark. Chips were then washed three times with DPBS before incubation with DAPI (1:5,000) for 5 min at room temperature in the dark, followed by two final DPBS washes prior to imaging.

### Confocal microscopy

Confocal images were acquired using a Zeiss LSM900 inverted laser scanning confocal microscope equipped with Airyscan 2 (Zeiss) and Plan-Apochromat 10× (Zeiss, NA = 0.45) and 20× (Zeiss, NA = 0.8) objective lenses. Imaging was performed using 405 nm, 488 nm, 561 nm and 640 nm laser lines.

### Statistical analysis

Data were analyzed using GraphPad Prism version 10 (La Jolla, CA). Statistical significance was assessed using unpaired two-tailed *t* tests (Fig. 1 and 4) or two-way analysis of variance (ANOVA) followed by Tukey’s or Šídák’s multiple-comparisons test (Fig. 2, 3, 5, 6), as detailed in the figure legends.

**Figure 1.**
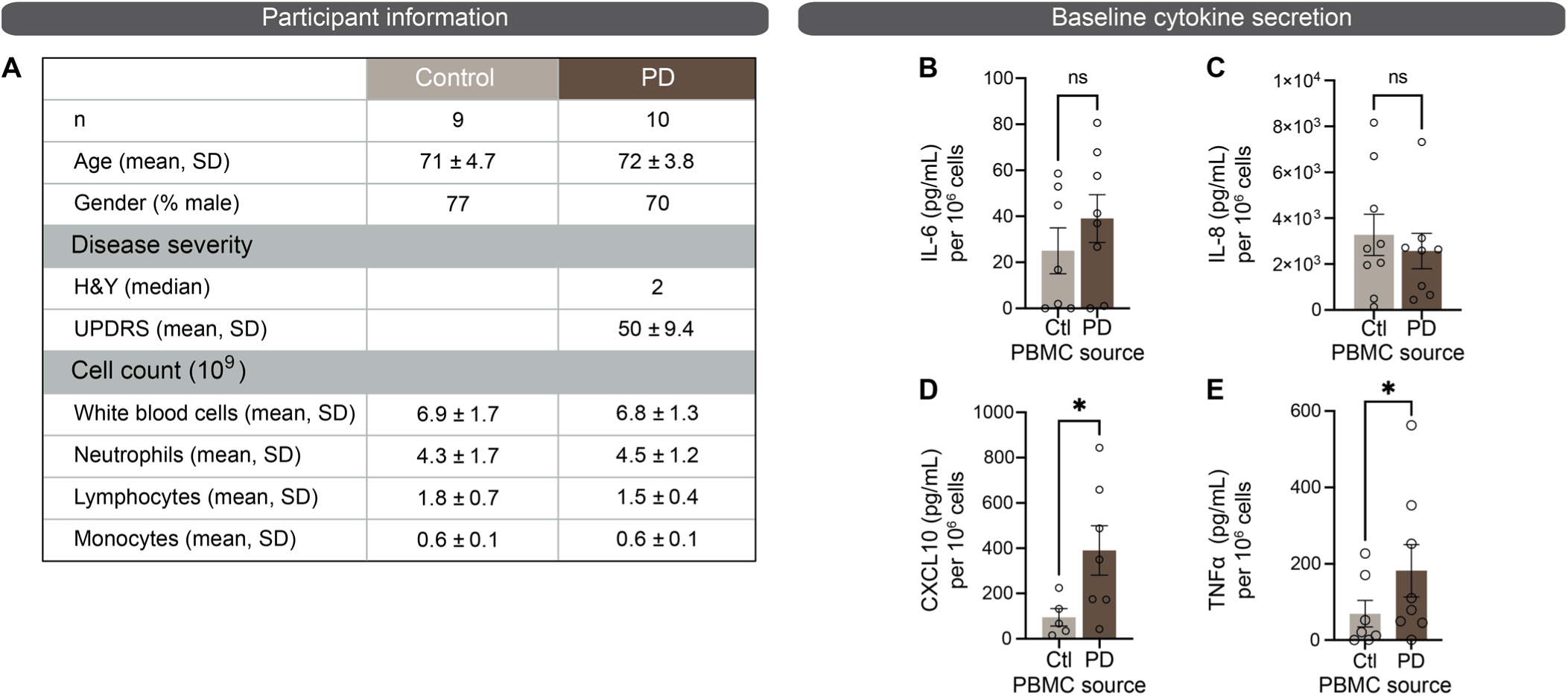
Baseline characteristics and cytokine secretion of control and PD PBMCs. (**A**) Baseline demographic and clinical characteristics of control and PD donors included in the study. (**B–E**) ELISA-based quantification of IL-6 (B), IL-8 (C), CXCL10 (D), and TNFα (E) in conditioned media collected from unstimulated PBMCs isolated from control and PD donors. Data are collected from 5 to 9 different PBMC donors; error bars represent mean ± SEM. Statistical analysis was performed using unpaired *t* test, *p≤0.05. Abbreviation: ns, not significant.

**Figure 2.**
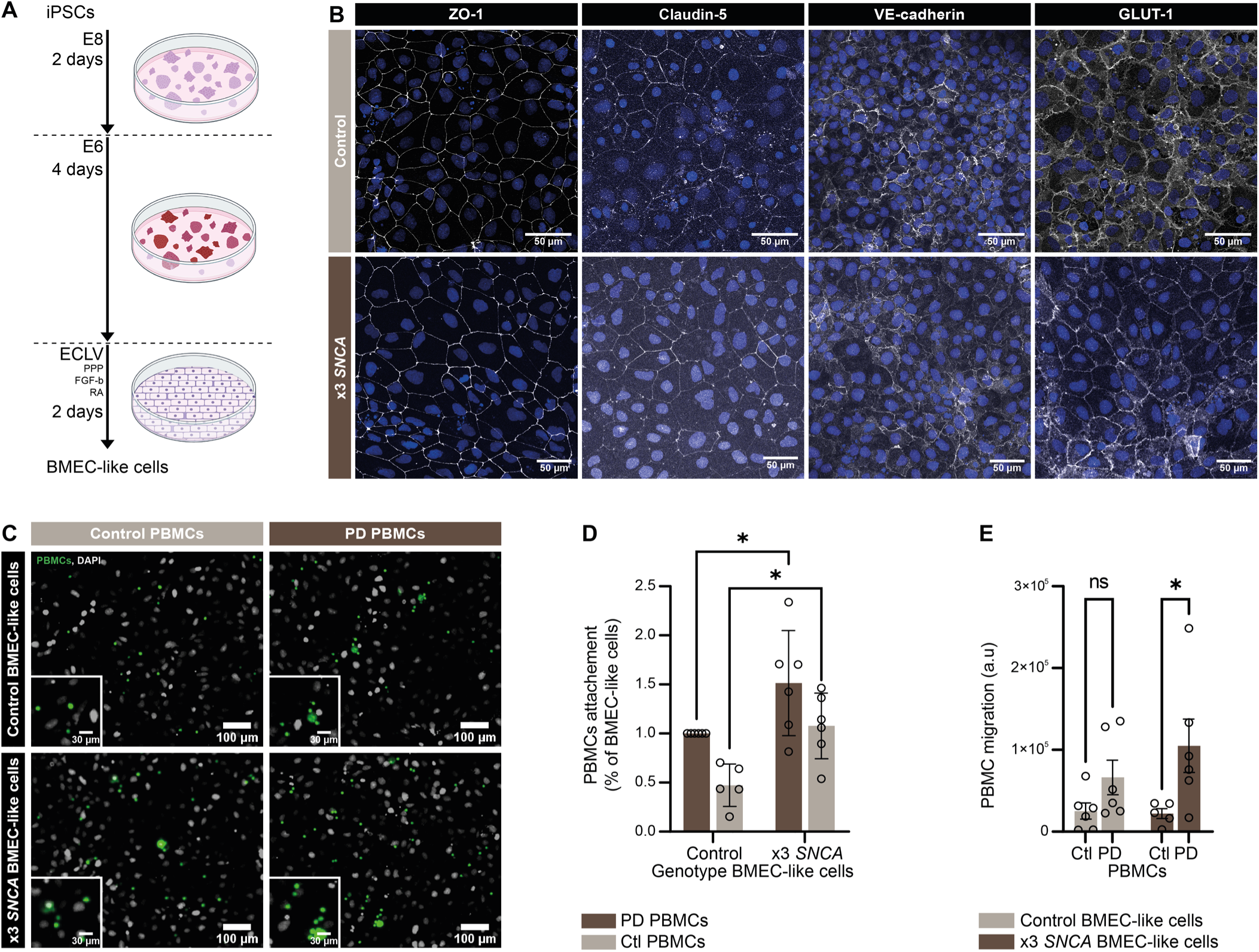
*SNCA* triplication enhances PBMC adhesion and transmigration in a static BBB model. (**A**) Schematic representation of the differentiation protocol used to generate iPSC-derived BMEC-like cells from control or *SNCA* triplication lines. (**B**) Confocal images of immunostained iPSC-derived BMEC-like cell monolayers generated from control or x3*SNCA* triplication iPSC lines, illustrating the expression of tight junction and endothelial markers anti-ZO-1, anti-Claudin-5, anti-VE-cadherin, and anti-GLUT1 (white). Nuclei are counterstained with DAPI (blue). Scale bars: 50 µm. (**C-D**) Representative images (C) and quantification (D) of PBMC attachment to control or x3*SNCA* BMEC-like cell monolayers following a 15 min incubation. (**E**) Quantification of PBMC transmigration across a static transwell BBB model. Error bars represent mean ± SEM. Statistical analysis was performed using two-way ANOVA followed by Šídák’s post-hoc test, *p≤0.05; Abbreviation: ns, not significant.

**Figure 3.**
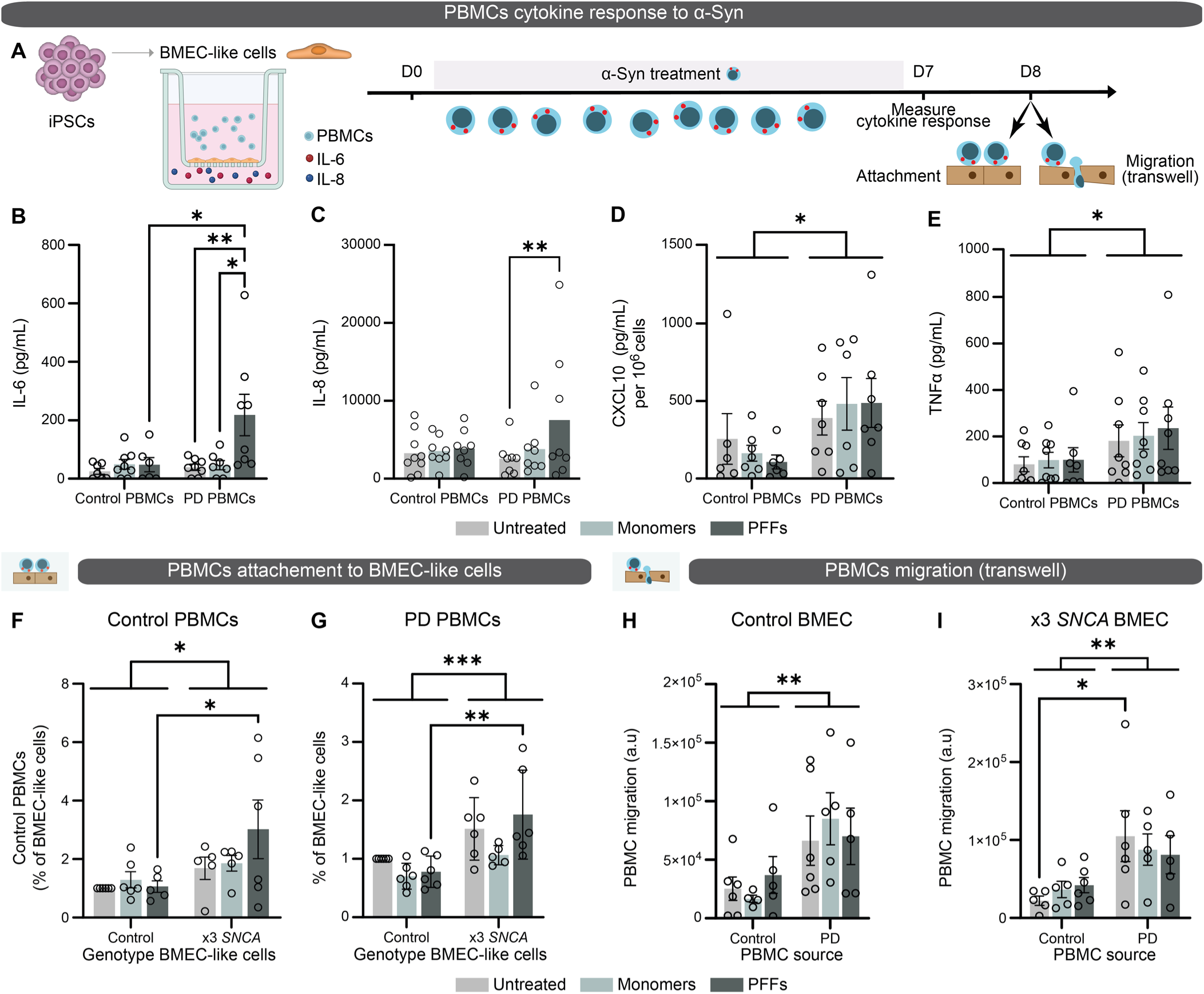
α-Syn pre-exposure modulates PBMC cytokine secretion and adhesion but not transmigration in a static model. (**A**) Schematic overview of the transwell model and experimental timeline used to assess the effects of α-Syn monomers or PFFs on PBMCs cytokine secretion, adhesion to BMEC-like cell monolayers, and transmigration in a static transwell BBB model. (**B–E**) ELISA-based quantification of IL-6 (B), IL-8 (C), CXCL10 (D), and TNFα (E) in conditioned media collected from control and PD PBMC cultures following exposure to vehicle control, α-Syn monomers (5 µg/mL), or PFFs (5 µg/mL) treatment. (**F–G**) Quantification of control (F) and PD (G) PBMCs adhesion to control or x3*SNCA* BMEC-like cell monolayers. (**H–I**) PBMCs transmigration across control (H) or x3*SNCA* (I) BMEC-like cell monolayers assessed using a static transwell BBB model. Data are from 5 to 6 independent experiments; error bars represent mean ± SEM. Statistical analysis were performed using two-way ANOVA followed by Tukey’s (B-E) or Šídák’s (F-I) post-hoc test, *p≤0.05, **p<0.01, ***p<0.001.

**Figure 4.**
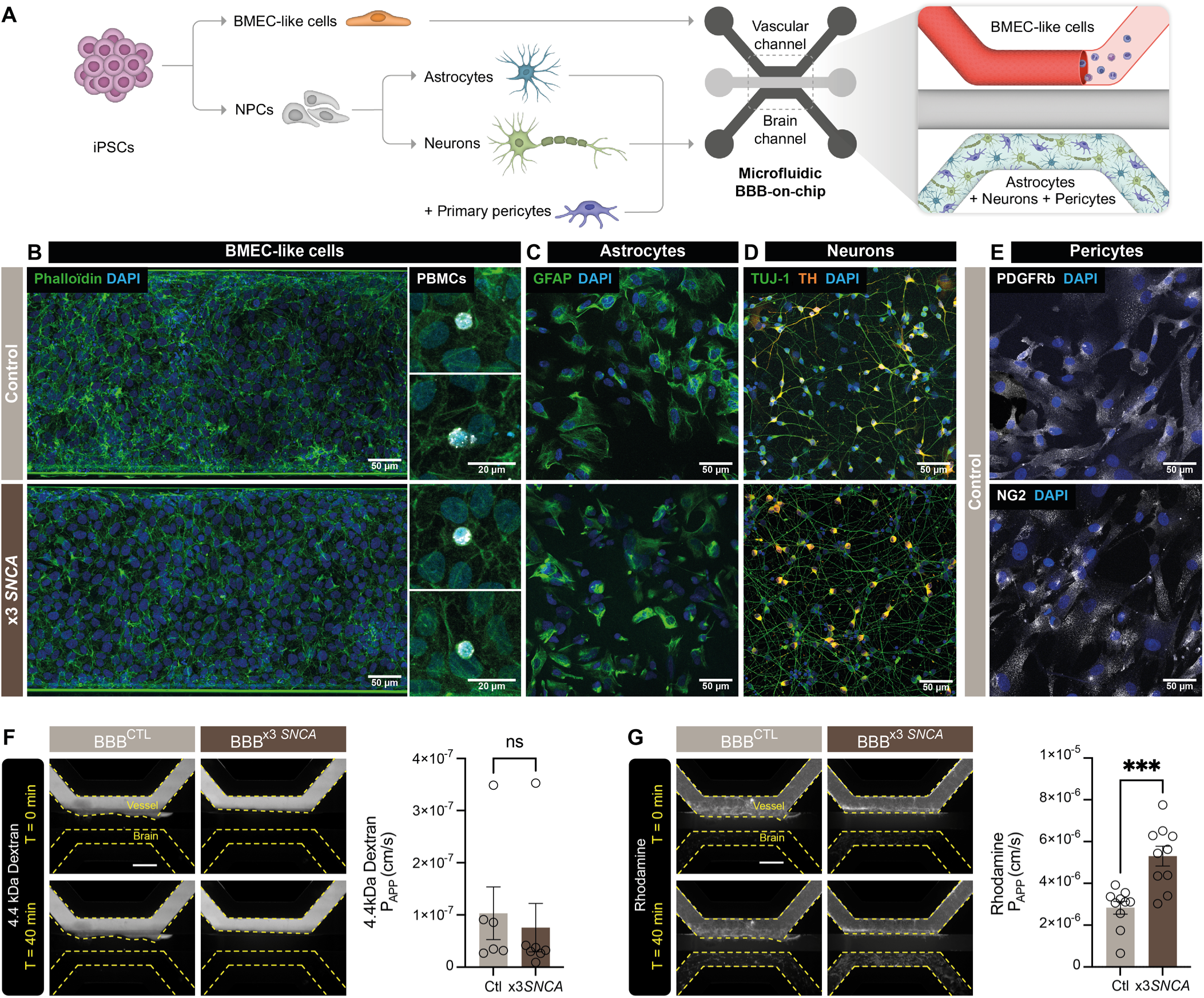
Engineering a microfluidic BBB chip incorporating x3*SNCA* mutation to model PD. (**A**) Illustration of the BBB chip model. (**B**) Confocal images of endothelial-like vessels located in the vascular compartment of the BBB chip. Actin filaments are visualized using phalloidin staining (orange), and the presence of peripheral immune cells is visualized using Cellbright labeling (white). (**C-E**) Confocal images of immunostained iPSC-derived cells confirming anti-GFAP expression (green) in astrocytes (C), anti-TUJ-1 (green) and anti-tyrosine hydroxylase (TH) (orange) in dopaminergic neurons (D), and anti-PDGFR-β (white) and anti-NG2 (white) in primary human pericytes (E). Nuclei are counterstained with DAPI (blue). Scale bars: 50 µm. (**F-G**) Representative fluorescence images showing the distribution of 3 kDa dextran (F) or rhodamine (G) in BBB^CTL^ and BBB^x3^ *^SNCA^* chips before and after a 40-min incubation with each fluorescent tracer. Graphs show quantification of apparent permeability (P_APP_) values at 4 days *in vitro.* Each data point represents P_APP_ measures of individual BBB chips of the Organoplate across 3 independent experiments. Scale bar: 600 µm (F, G). Data are represented as mean ± SEM. Statistical analysis was performed using an unpaired two-tailed t-test, ***p<0.001; Abbreviation: ns, not significant.

**Figure 5.**
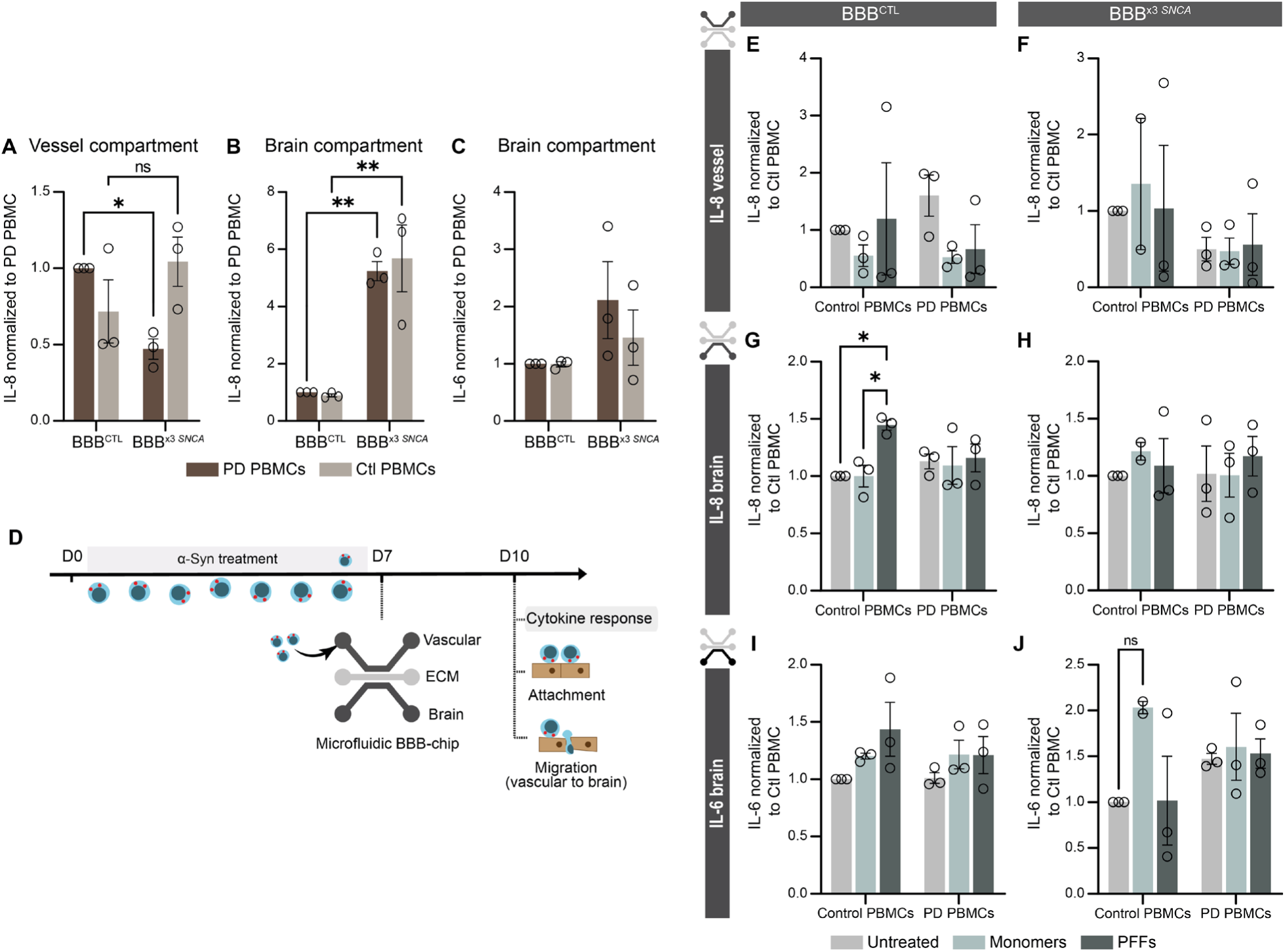
Cytokine secretion in vascular and brain compartments of the microfluidic BBB chip. **(A-C)** ELISA-based quantification of basal IL-8 (A-B) and IL-6 (C) levels in conditioned media collected from the brain compartment of BBB^CTL^ and BBB^x3^ *^SNCA^* chips following perfusion with unstimulated control or PD PBMCs. **(D)** Schematic representation of the experimental design used to assess PBMCs attachment and transmigration in the BBB chip following pre-exposure to vehicle control, α-Syn monomers (5 µg/mL) or PFFs (5 µg/mL). PBMCs were treated for 7 days prior to perfusion into the vascular compartment of the BBB chip. (**E-J**) ELISA-based quantification of cytokine levels in conditioned media collected from the vascular or brain compartments of BBB^CTL^ and BBB^x3^ *^SNCA^* chips following perfusion with control or PD PBMCs pre-exposed to vehicle control, α-Syn monomers (5 µg/mL), or PFFs (5 µg/mL). IL-8 levels were quantified in the vascular compartment of BBB^CTL^ (E) and BBB^x3^ *^SNCA^* (F). IL-8 and IL-6 levels were also quantified in the brain compartment of BBB^CTL^ (G, I) and BBB^x3^ *^SNCA^* chips (H, J). Data are from 3 independent biological replicates; error bars represent mean ± SEM. Statistical analyses were performed using two-way ANOVA followed by a Šídák’s (A-C) or Tukey’s (E-J) post-hoc test, *p≤0.05, **p<0.01; Abbreviation: ns, not significant.

**Figure 6.**
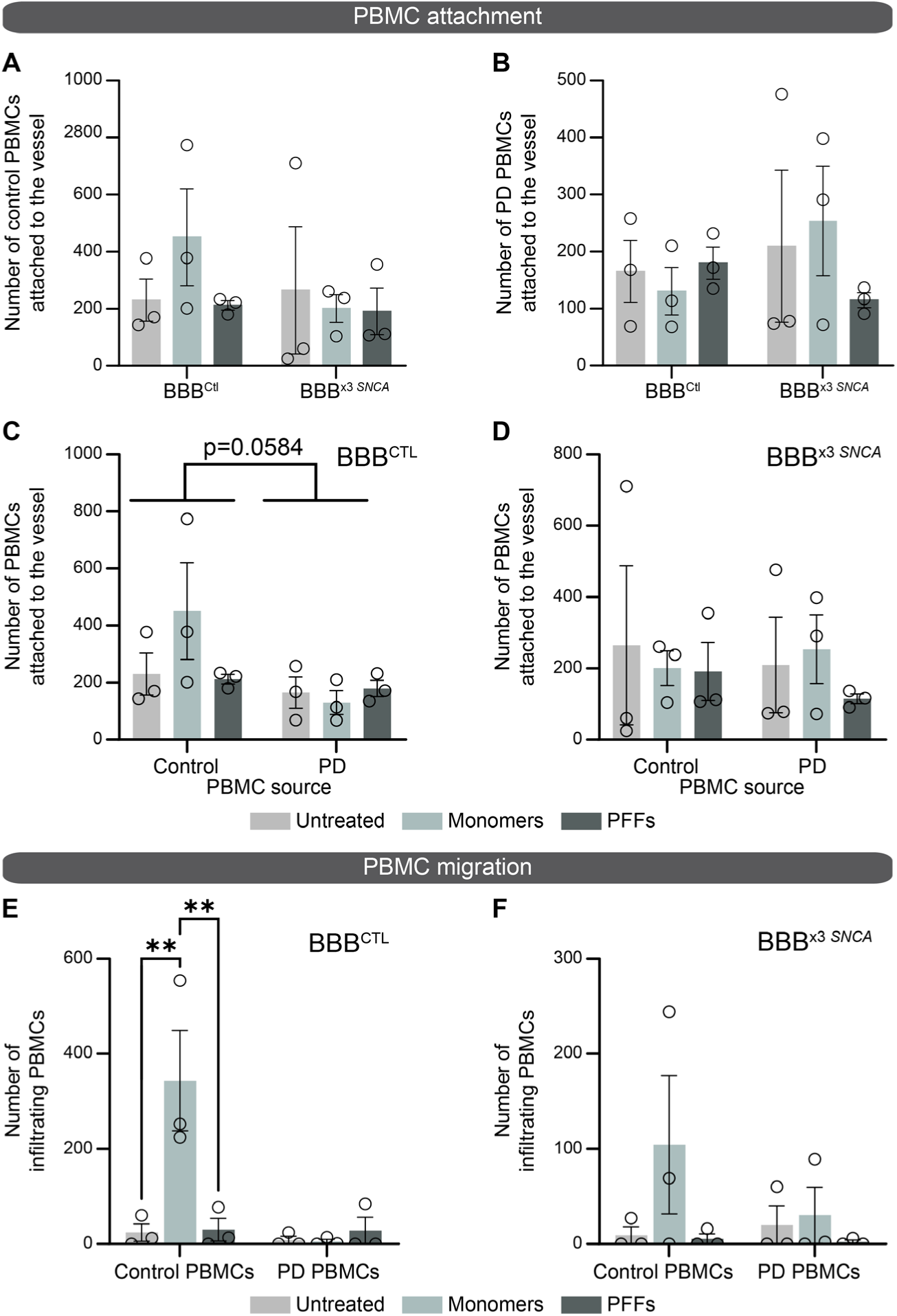
PBMCs adhesion and transmigration in the microfluidic BBB chip following α-Syn pre-exposure. (**A-B**) Quantification of PBMC attachment to endothelial-like vessels. The number of control (A) or PD (B) PBMCs attached to endothelial-like vessels was quantified in BBB^CTL^ and BBB^x3^ *^SNCA^* chips. (**C-D**) Alternative representation of PBMCs attachment data from panels A-B, presented here enable direct statistical comparison of BBB genotype-related effects. (**E-F**) Quantification of PBMC transmigration from the vascular to the brain compartment in BBB^CTL^ (E) or BBB^x3^ *^SNCA^*(F) chips. Data are from 3 independent experiments; error bars represent mean ± SEM. Statistical analyses were performed using two-way ANOVA followed by Tukey’s post-hoc test, **p<0.01.

## Results

### Participant characteristics and baseline cytokine secretion

Human PBMCs were obtained from 9 control donors (77% males; mean age, 71 years) and 10 individuals diagnosed with PD (70% males; mean age, 72 years). Control and PD cohorts were comparable with respect to age, sex distribution, and circulating immune cell counts (**Fig. 1A**). PD participants presented a median Hoehn and Yahr (H&Y) score of 2 and a mean Unified PD Rating Scale (UPDRS) score of 50 ± 9.4, consistent with mild-to-moderate disease severity. Following blood collection, PBMCs were isolated and analyzed to assess baseline immune cell composition. PBMC preparations from control donors contained an average of 6.9×10^9^ white blood cells, consisting of an average of 4.3×10^9^ neutrophils, 1.8×10^9^ lymphocytes, and 6×10^8^ monocytes. Similarly, PBMC samples from PD patients consisted of an average of 6.8×10^9^ white blood cells, comprising 4.5×10^9^ neutrophils, 1.5×10^9^ lymphocytes and 6×10^8^ monocytes, indicating comparable baseline immune cell distributions between groups. Baseline cytokine secretion by resting PBMCs revealed similar levels of IL-6 and IL-8 in control and PD-derived samples (**Fig. 1B-C**). In contrast, significant differences were observed for CXCL10 and TNF-α secretion (**Fig. 1D-E**). Control PBMCs exhibited a mean baseline secretion of 100 pg/mL CXCL10, whereas PBMCs derived from PD patients displayed significantly higher levels, averaging approximately 400 pg/mL. Likewise, TNF-α secretion increased from approximately 50 pg/mL in control PBMCs to nearly 200 pg/mL in PBMCs isolated from PD donors.

### *SNCA* triplication promotes PBMC-BMEC interactions in the static BBB model

Control and PD PBMCs were next assessed for their ability to interact with monolayers of iPSC-derived BMEC-like cells generated from control donors (BMEC^CTL^) or PD patients harboring the *SNCA* triplication mutation (BMEC^x3*SNCA*^, **Fig. 2A**). Successful differentiation of iPSCs into BMEC-like cells was confirmed by immunofluorescence-based detection of the endothelial and tight junction markers ZO-1, claudin-5, VE-cadherin, and ZO-1 (**Fig. 2B**). Fluorescently labelled PBMCs were incubated on BMEC-like cell monolayers for 15 min, followed by gentle washing to remove non-adherent cells, and PBMC attachment was quantified by fluorescence microscopy. Both control and PD PBMCs exhibited significantly increased attachment to BMEC^x3*SNCA*^ compared with BMEC^CTL^ monolayers. PBMCs attachment to the endothelial layer increased by at least 50% when the BMEC genotype harbored the *SNCA* triplication mutation, irrespective of PBMC donor status (**Fig. 2C-D**). PBMCs transmigration was assessed using a static transwell BBB model in which BMEC^CTL^ or BMEC^x3*SNCA*^ were cultured on membrane inserts forming the luminal compartment, while the abluminal compartment was supplemented with IL-6 and IL-8 (50 ng/mL each) to establish a chemotactic gradient. Under these conditions, no significant differences in transmigration were observed between control and PD PBMCs BMEC^CTL^ monolayers, whereas PD-derived PBMCs exhibited significantly increased transmigration compared with control PBMCs across BMEC^x3*SNCA*^ monolayers (**Fig. 2E**).

### α-Syn PFFs modulate PBMC attachment but not transmigration in the static BBB model

The effect of α-Syn species on PBMC behavior was next evaluated by pre-exposing control and PD PBMCs to α-Syn monomers and PFFs for 7 days, followed by analysis of cytokine secretion, attachment, and transmigration in static BBB models (**Fig. 3A**). Following α-Syn pre-exposure, changes in cytokine secretion were observed selectively in PD-derived PBMCs. Specifically, treatment with PFFs, but not monomers, resulted in a significant increase in IL-6 and IL-8 secretion by PD PBMCs compared with untreated cells, from ∼40 to 200 pg/mL and ∼2,500 to 8,000 pg/mL, respectively (**Fig. 3B-C**). In contrast, CXCL10 and TNF-α secretion remained consistently higher in PD PBMCs compared with control PBMCs, irrespective of α-Syn treatment (**Fig. 3D-E**). PBMC attachment to BMEC-like cell monolayers was subsequently assessed. Exposure to PFFs significantly increased the attachment of both control and PD PBMCs to BMEC^CTL^ and BMEC^x3*SNCA*^ monolayers, whereas α-Syn monomers had no detectable effect (**Fig. 3F-G**). Finally, PBMCs transmigration was evaluated using the static transwell BBB model. Neither α-Syn monomers nor PFFs treatment significantly altered PBMCs migration. Instead, migration was primarily determined by donor PBMC disease status, with PD-derived PBMCs exhibiting increased transmigration compared with control PBMCs (**Fig. 3H-I**).

### Microfluidic BBB chip modeling of PD-related *SNCA* triplication mutation

PBMC behavior was next evaluated in a 3D microfluidic BBB chip model, in which BMEC-like cells formed a perfusable 3D endothelial vessel (**Fig. 4A**). The vascular compartment containing circulating PBMCs was visualized by confocal microscopy (**Fig. 4B**), and the adjacent CNS compartment consisted of iPSC-derived astrocytes, dopaminergic neurons, and human primary pericytes (**Fig. 4A, C-E**, **Suppl. Fig. 1**). Fluid shear stress was induced by the regular movement of a plate rocker (7° angle every 8 min), which promoted vessel formation and ensured vascular perfusion. This platform achieved an estimated peak shear stress of 1.2 dyne/cm² and a peak flow rate of approximately 13.2 μL/min ^27^. Under these conditions, we perfused 45,000 PBMCs per vessel to deliver ∼5.0 × 10⁴ PBMCs/min/cm² and reproduce physiological *in vivo* cerebral PBMC flux range. To ensure functional integrity of the endothelial vessel, barrier integrity assays were performed prior to introducing PBMCs (**Fig. 4F-G**). Data showed low apparent permeability (P_AAP_) values when a 3 kDa tracer was perfused in BBB^CTL^ and BBB^x3*SNCA*^, suggesting functional integrity of endothelial tight junctions in both models. In contrast, P_AAP_ quantification of the p-glycoprotein substrate rhodamine revealed increased vascular-to-brain diffusion in BBB^x3*SNCA*^ compared to BBB^CTL^, suggesting reduced substrate efflux in the PD model.

### The BBB chip model is immuno-responsive

Cytokine secretion was quantified in the vascular and CNS compartments of the microfluidic chips to assess the immune competence of the model. Unstimulated control and PD PBMCs were incubated in the vascular compartment of BBB^CTL^ or BBB^x3^ *^SNCA^* chips for 72h, and ELISA-based quantification confirmed IL-8 secretion in the vascular compartment. Unstimulated PD PBMCs reduced ∼2-fold IL-8 secretion in BBB^x3*SNCA*^ vessels compared to BBB^CTL^, while this effect was not observed in vessels incubated with control PBMCs (**Fig. 5A**), indicating that PBMC disease status is sufficient to modulate vascular responses independently of α-Syn treatment. In parallel, conditioned media collected from the CNS compartment revealed that glial cells secreted both IL-8 and IL-6 (**Fig. 5B-C**), with BBB^x3*SNCA*^ chips displaying nearly 5-fold higher IL-8 secretion compared to BBB^CTL^, regardless of PBMC disease status (**Fig. 5B**). Then, to assess the effect of α-Syn treatment, control and PD PBMCs were pre-stimulated with 5 μg/mL α-Syn monomers, PFFs, or PBS vehicle for a total of 7 days, and incubated in the vascular compartment of BBB^CTL^ or BBB^x3*SNCA*^ chips for 72 h (**Fig. 5D**). Data suggest that vascular IL-8 secretion remained unchanged across α-Syn treatment conditions (**Fig. 5E-F**). However, analysis of conditioned media from the CNS compartment revealed that control (but not PD) PBMCs exposed to PFFs triggered an increased secretion of IL-8 in the BBB^CTL^ brain (**Fig. 5G-H**), but IL-6 secretion did not significantly differ (**Fig. 5I-J**).

### α-Syn monomers, not PFFs, increase PBMCs transmigration in the BBB chip model

Next, we mirrored the PBMC attachment and infiltration experiments performed in the monolayer and transwell platforms. The number of PBMCs attached to the 3D BMEC-like vessel was counted, and there was no significant difference in cell attachment induced by α-Syn treatment or BMEC-like cell genotype. However, when incubated in the control BBB chip, PD PBMCs attached less readily to the endothelial-like vessel compared to PBMCs derived from control donors (**Fig. 6A-D**). This observation suggests that, in the BBB chip, PBMC disease status affects cell attachment, while in the monolayer, pre-exposure to α-Syn is the strongest modifier of immune cell-endothelium interaction. Then, the measure of PBMC infiltration from the vascular to the CNS compartment of the BBB chip further revealed differences between the transwell and microfluidic model. In contrast to the transwell, PBMC transmigration in the microfluidic BBB was significantly increased by α-Syn pre-treatment, and particularly pre-exposure to α-Syn monomers (**Fig. 6E-F**). Here, data show a predominant response of control vs PD PBMCs, which contrasts with the transwell assay.

## Discussion

### PD-associated immune cells exhibit altered interactions with the BBB

This study aimed to assess PBMC-BBB interactions in a transwell vs. 3D microfluidic BBB chip to support studies on the peripheral-brain immune axis in neurodegenerative diseases, and in particular PD. These two models were generated using human iPSCs, and they were exposed to PBMCs isolated from the blood of control donors or people with PD at a moderate disease stage (H&Y=2). A comparative analysis of PBMC attachment and transmigration in the transwell and microfluidic BBB models confirmed that peripheral immune cells isolated from the blood of people with PD interact differently with BMEC-like cells compared to control PBMCs. Furthermore, the genotype of BMEC-like cells and pre-exposure to α-Syn PFF altered immune-endothelial interactions, emphasizing that cell-cell communication at the brain endothelium is a multi-component, dynamic process that can be affected by PD-related risk factors. The observation that both endogenous α-Syn overexpression and exogenous PFF exposure altered PBMC trafficking suggests that multiple α-Syn-related mechanisms may converge on common pathways regulating BBB-immune interactions. One possibility is that extracellular α-Syn may act at the cell surface by engaging pattern-recognition or adhesion-related receptors expressed on PBMCs, thereby modifying inflammatory signaling and cell-endothelial interactions. Previous studies demonstrated that α-Syn can engage TLR1/2 and CD11b on microglia ^37–39^, and similar mechanisms may occur when peripheral immune cells encounter pathological α-Syn assemblies in the systemic circulation. Alternatively, other studies have shown rapid uptake of α-Syn by endothelial and immune cells, leading to intracellular signaling responses ^40,41^. These mechanisms are not mutually exclusive and may operate simultaneously. As α-Syn uptake was not assessed in the present study, future work will be needed to determine the relative contributions of cell-surface signaling and intracellular α-Syn processing to the observed effects.

### Complementary BBB models of α-Syn pathology identify dysregulated immune-endothelial interactions

Each BBB model captured a different aspect of PBMC-BMEC interactions. In the 2D transwell system, *SNCA* triplication increased PBMC attachment to endothelial-like monolayers, and exposure to α-Syn PFFs further enhanced attachment. In contrast, the 3D microfluidic model showed reduced PBMC attachment in PD samples compared with controls, and α-Syn PFFs did not significantly affect attachment. Regarding transmigration, the 2D model showed increased migration of PD PBMCs compared with controls, independent of endothelial genotype or α-Syn exposure. This effect was not recapitulated in the 3D BBB chip, where α-Syn monomer pre-treatment instead increased transmigration of control PBMCs from the vascular to the parenchymal compartment. Importantly, the 3D BBB model was immune-competent, as both endothelial-like and glial cells altered cytokine secretion in response to PBMC exposure, consistent with previous validation of inflammatory responsiveness ^29^. PBMCs from individuals with PD also displayed a dysregulated immune phenotype consistent with previous reports and multi-omics studies demonstrating systemic immune dysregulation in PD^42–47^. Published studies have also reported that PD PBMCs display altered secretory responses to pathological α-Syn exposure *ex-vivo* ^48,49^ and, together with data provided in this study, these observations support the concept that circulating immune cells in PD exist in a distinct activation state that may influence their interactions with the cerebral vasculature in 2D and 3D models. Our findings that PD PBMCs display altered BBB trafficking behavior therefore aligns with growing evidence that peripheral immune dysfunction represents an integral component of PD pathophysiology.

### Model architecture influences PBMC-BBB interactions

Overall, both models indicated PD-relevant alterations in PBMC-endothelial interactions. However, the dimensionality and architecture of the systems strongly influenced the observed cellular behavior. The microfluidic BBB chip more closely reflects physiological exposure conditions at the human BBB, whereas the static transwell system imposes prolonged immune-endothelial contact. A previously published study compared static and dynamic BBB models built using immortalized hCMEC/D3 cells and human primary astrocytes, and measures of transendothelial electrical resistance (TEER) and P_AAP_ reflected increased barrier integrity in microfluidic devices compared to transwell systems ^50^. In addition, the migration of healthy PBMCs across hCMEC/D3 cells was reduced by ∼75% in dynamic vs. static flow conditions. Therefore, the dynamic physiological environment and fluid flow characteristic of microfluidic systems promotes barrier function, and influences PBMC dynamics by altering the timing and duration of endothelial contacts compared with static conditions. Shear stress is known to regulate endothelial phenotype, glycocalyx organization, barrier properties, and the expression of adhesion molecules involved in immune cell trafficking ^51–54^, processes that are absent in static culture systems. Moreover, physiological flow supports the sequential recruitment cascade of circulating immune cells, including transient rolling, activation, firm adhesion, and subsequent transmigration, which may differentially shape immune cell behavior compared with static conditions. These flow-dependent properties highlight fluid dynamics as a key regulator of endothelial responses and support the use of dynamic BBB models to study immune-endothelial interactions under physiologically relevant conditions ^55–57^. Static and dynamic platforms differ in geometry, compartmentalization, and culture conditions, all of which influence experimental readouts and require careful interpretation ^16,58–60^. Accordingly, model-specific constraints should be considered when comparing results across systems. Future studies will be needed to determine which adhesion pathways and immune cell subsets contribute to the distinct trafficking profiles observed across BBB platforms. Together, these findings demonstrate that PBMC-BBB interactions in PD are shaped by immune cell status, endothelial genotype, and α-Syn exposure, but are strongly modulated by BBB model dimensionality and flow conditions.

Several limitations should be acknowledged. First, the number of donor-derived PBMC samples was limited, reflecting the challenges associated with recruiting clinically characterized patients and performing complex BBB chip experiments. Although consistent trends were observed across independent experiments, larger cohorts will be required to validate the robustness and generalizability of these findings. Second, PBMCs were analyzed as a heterogeneous population, precluding identification of the specific immune subsets driving the observed phenotypes. Future studies incorporating immune phenotyping by flow cytometry and/or single-cell transcriptomics will be important to define the relative contributions of monocytes, T cells, B cells, and other circulating immune populations to BBB interactions and transmigration behavior. In addition, integrating additional inflammatory or genetic PD-relevant triggers (e.g. environmental toxins) would further refine these models and help establish standardized benchmarks for reproducible neurovascular research.

In conclusion, this study provides a controlled human platform to evaluate how immune phenotypes are differentially captured across BBB model systems, thus supporting follow-up studies to define the molecular mechanisms underlying PBMC dysregulation in PD. This study underscores the importance of incorporating physiologically relevant multicellular and dynamic BBB systems to more accurately capture neuroimmune interactions. More broadly, the complementary use of static and dynamic models provides a framework to assess the robustness and model-dependence of experimental readouts in BBB research. The choice of model system, including considerations of flow, cellular composition, and throughput, should therefore be guided by the specific experimental questions and desired physiological relevance.

## Acknowledgements

The authors would like to express their gratitude to the people who donated blood samples, and therefore enabled the completion of this study. Funding was provided by the Michael J Fox Foundation (to AdRJ and FC), the Parkinson’s Foundation (Launch Award), the Fondation du CHU de Québec, and a Fond de Recherche du Québec en Santé (FRQS) Junior 1 Research Scholar Award to AdRJ. OA was supported by graduate student fellowships from Desjardins and Fondation du CHU de Québec, and Parkinson Canada. FB was supported by a postdoctoral fellowship from Parkinson Canada. EB is recipient of a Merite Award from the FRQS.

## Authors contributions (using CRediT taxonomy, available at https://casrai.org/credit)

OM: Investigation, Visualization, Writing—original draft.; LG: Investigation, Visualization, Writing—original draft.; FB: Visualization, Writing—review & editing; AY: Investigation; OA: Investigation; FNAMB: Investigation; MD: Investigation; KL: Supervision, Resources, Writing—review & editing; FC: Supervision, Funding acquisition, Writing—review & editing; AdRJ: Conceptualization, Investigation, Formal Analysis, Methodology, Validation, Visualization, Supervision, Funding acquisition, Writing—original draft.

---

**Supplementary Figure 1.**
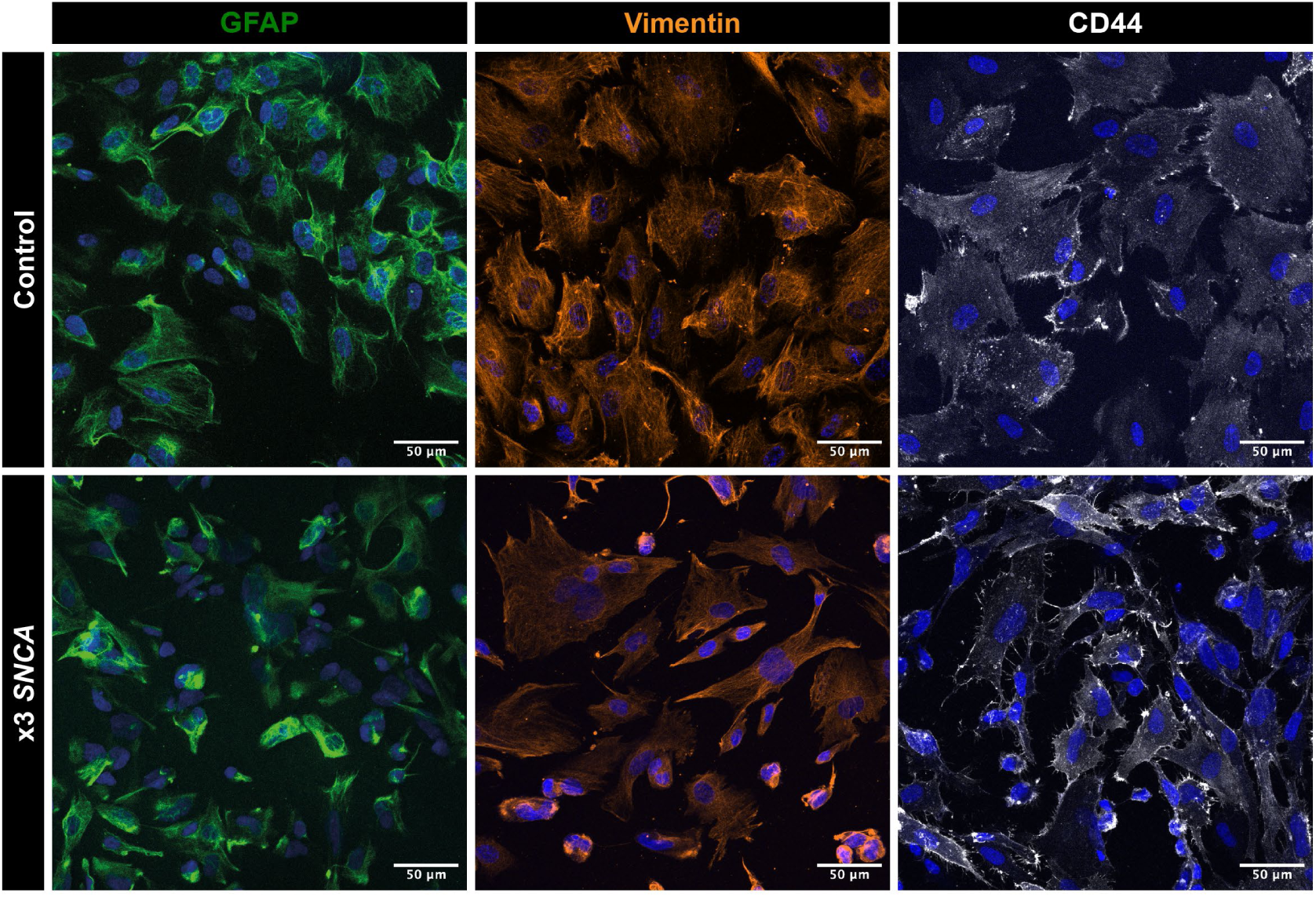
iPSC-derived astrocytes express cell type-relevant markers. Confocal images of immunostained iPSC-derived astrocytes showing anti-GFAP (green), anti-Vimentin (orange) and anti-CD44 (white). Nuclei are counterstained with DAPI (blue). Astrocyte quality assessment reflects previously published reports ^29,31^.

